# Enhancing KCC2 activity decreases hyperreflexia and spasticity after chronic SCI

**DOI:** 10.1101/2020.04.25.061176

**Authors:** Jadwiga N. Bilchak, Kyle Yeakle, Guillaume Caron, Dillon C. Malloy, Marie-Pascale Côté

**Affiliations:** Marion Murray Spinal Cord Injury Research Center, Department of Neurobiology and Anatomy, Drexel University College of Medicine, Philadelphia, PA 19129

**Author notes:** Corresponding author Marie-Pascale Côté, Ph.D. (corresponding author), Assistant Professor, Drexel University College of Medicine, Department of Neurobiology and Anatomy, Philadelphia, PA 19129.

**Keywords:** spinal cord injury, rehabilitation, chloride homeostasis, neuroplasticity, KCC2, CLP257

## Abstract

After spinal cord injury (SCI), the majority of individuals develop spasticity, a debilitating condition involving involuntary movements, co-contraction of antagonistic muscles, and hyperreflexia. By acting on GABAergic and Ca^2+^-dependent signaling, current anti-spastic medications lead to serious side effects, including a drastic decrease in motoneuronal excitability which impairs motor function and rehabilitation efforts. Exercise, in contrast, decreases spastic symptoms without decreasing motoneuron excitability. These functional improvements coincide with an increase in expression of the chloride co-transporter KCC2 in lumbar motoneurons. Thus, we hypothesized that spastic symptoms can be alleviated directly through restoration of chloride homeostasis and endogenous inhibition by increasing KCC2 activity. Here, we used the recently developed KCC2 enhancer, CLP257, to evaluate the effects of acutely increasing KCC2 extrusion capability on spastic symptoms after chronic SCI. Sprague Dawley rats received a spinal cord transection at T12 and were either bike-trained or remained sedentary for 5 weeks. Increasing KCC2 activity in the lumbar enlargement improved the rate-dependent depression of the H-reflex and reduced both phasic and tonic EMG responses to muscle stretch in sedentary animals after chronic SCI. Furthermore, the improvements due to this pharmacological treatment mirror those of exercise. Together, our results suggest that pharmacologically increasing KCC2 activity is a promising approach to decrease spastic symptoms in individuals with SCI. By acting to directly to restore endogenous inhibition, this strategy has potential to avoid severe side effects and improve the quality of life of affected individuals.

**Significance Statement:** Spasticity is a condition that develops after spinal cord injury (SCI) and causes major complications for individuals. We have previously reported that exercise attenuates spastic symptoms after SCI through an increase in expression of the chloride co-transporter KCC2, suggesting that restoring chloride homeostasis contributes to alleviating spasticity. However, the early implementation of rehabilitation programs in the clinic is often problematic due to co-morbidities. Here, we demonstrate that pharmacologically enhancing KCC2 activity after chronic SCI reduces multiple signs of spasticity, without the need for rehabilitation.

## Introduction

Disruption in synaptic inhibition is a common feature of an ever-growing list of diseases, ranging from neurodevelopmental conditions such as Down syndrome, epilepsy, and autism, to psychiatric disorders including schizophrenia and depression (De Koninck, 2007; Ben-Ari et al., 2012; Kaila et al., 2014; Ben-Ari, 2017). Targeting chloride equilibrium has therefore become recognized as a promising therapeutic strategy (De Koninck, 2007; Ben-Ari et al., 2012; Gagnon et al., 2013; Kahle et al., 2014; Puskarjov et al., 2014). Recently, the critical importance of chloride homeostasis in the context of spinal cord injury (SCI) has been acknowledged with the identification of its involvement in spasticity and neuropathic pain (see also Coull et al., 2003; Boulenguez et al., 2010; Sanchez-Brualla et al., 2018; Mapplebeck et al., 2019; reviewed in Côté, 2020).

After SCI, there is a progressive decrease in expression of the chloride extruder KCC2 (Boulenguez et al., 2010) that plateaus 4 weeks post-injury (Côté et al., 2014). The subsequent decrease in chloride extrusion drives the system toward a state resembling early development, in which GABA_A_-mediated responses are depolarizing (Payne et al., 2003). Thus, after SCI there is reduced inhibition, decreased ability to suppress excitatory events, and facilitation of incoming excitatory inputs (Hubner et al., 2001; Jean-Xavier et al., 2006; Vinay and Jean-Xavier, 2008; Boulenguez et al., 2010; Bos et al., 2013). In adult lumbar motoneurons, the disruption in chloride homeostasis has been associated with the development of spasticity (Vinay and Jean-Xavier, 2008; Boulenguez et al., 2010; Viemari et al., 2011; Bos et al., 2013). Spasticity is characterized by a velocity-dependent increase in the stretch reflex which leads to symptoms such as increased muscle tone, hyperreflexia, painful muscle contractures, and co-contraction of antagonist muscles (Nielsen et al., 2007). This debilitating condition emerges in up to 75% of SCI individuals, and critically hampers functional recovery (Maynard et al., 1990; Skold et al., 1999; Biering-Sorensen et al., 2006; Holtz et al., 2017).

Most anti-spastic drugs currently available act upstream of GABA and Ca^2+^-dependent mechanisms, leading to serious side effects. These include a deep depression of CNS excitability, significant reductions in muscle activity, and muscle weakness, all of which further impede residual motor function and its recovery (Dario and Tomei, 2004; Taricco et al., 2006; Lapeyre et al., 2010; Simon and Yelnik, 2010; Angeli et al., 2012). There is a great need for strategies to combat spasticity that avoid depressing CNS excitability and muscle activity. While activity-based therapies are effective in a subset of SCI individuals (Petropoulou et al., 2007; Elbasiouny et al., 2010; Dietz and Sinkjaer, 2012), they are expensive and not widely available. In addition, SCI is accompanied by many co-morbidities that make exercise difficult or even impossible for some patients (Yelnik et al., 2009; Simon and Yelnik, 2010), especially early after injury. We have previously shown that exercise prevents the development of spasticity by increasing KCC2 expression (Côté et al., 2014; Beverungen et al., 2019). Thus, a possible alternative to current anti-spastic drugs and to activity-based therapies is to restore chloride homeostasis by increasing KCC2.

Recently, a family of selective KCC2 activators known as CLPs was developed, allowing us to directly target KCC2 activity (Gagnon et al., 2013; Kahle et al., 2014; Ferrini et al., 2017). CLPs have been shown to restore impaired KCC2-mediated Cl^−^ extrusion in neurons *in vitro* and *in vivo* (Gagnon et al., 2013), and were later shown to increase KCC2 activity in various disease models in which KCC2 is pathologically decreased (Gagnon et al., 2013; Ostroumov et al., 2016; Ferrini et al., 2017; Chen et al., 2018; Thomas et al., 2018; Lizhnyak et al., 2019).

Here, we show that pharmacologically increasing KCC2 activity in chronic SCI animals with CLP257 reduces hyperreflexia and spastic symptoms. Furthermore, we demonstrate that this pharmacological treatment mimics the beneficial effects of rehabilitation. This work paves the way for future investigation of KCC2 enhancers as alternatives to exercise-based therapies for the treatment of spasticity after SCI.

## Materials & Methods

### Experimental Design

In a rat model of complete thoracic SCI (T12), we investigated the effects of pharmacologically increasing KCC2 activity on spasticity and hyperreflexia and how this pharmacological approach interacts with exercise. The KCC2 enhancer, CLP257, was used to increase KCC2 activity in chronic SCI rats, whereas Control rats received saline. SCI animals were randomly assigned to one of the following groups: chronic spinal cord injured + CLP257 (SCI + CLP257; n=10), chronic spinal cord injured + saline (SCI; n= 8), chronic spinal cord injured + bike-training + CLP257 (SCI + Ex + CLP257; n=11), and chronic spinal cord injured + bike-training + saline (SCI + Ex; n=5). Five weeks post injury, a terminal experiment was conducted, in which CLP257 or saline was administered to the relevant groups. The effect of restoring chloride homeostasis on hyperreflexia and spasticity was assessed by measuring the excitability of the H-reflex and the rate-dependent depression, as well as muscle forces and EMG activity in response to ramp-hold-release stretches, before and after application of either CLP257 or saline.

All procedures were performed in accordance with protocols approved by Drexel University College of Medicine Institutional Animal Care and Use Committee, followed National Institutes of Health guidelines for the care and use of laboratory animals, and complied with Animal Research: Reporting of In Vivo Experiments (ARRIVE).

### Surgical procedures and postoperative care

Adult female Sprague Dawley rats (240-300g, Charles River Laboratories) underwent a complete spinal transection at the low thoracic level (T12) as described previously (Côté et al., 2011; Côté et al., 2014; Beverungen et al., 2019). Briefly, rats were anesthetized with isoflurane (1-4%) in O_2_ and, under aseptic conditions, a laminectomy was performed at the T10–T11 vertebral level. The dura was carefully slit open, the spinal cord completely severed with small scissors, and the cavity filled with absorbable hemostats (Pfizer, New York, NY, USA) to promote homeostasis. The completeness of the lesion was ensured by the distinctive retraction of the rostral and caudal spinal tissue and by examining the ventral floor of the spinal canal during surgery, and was later confirmed post-mortem. Paravertebral muscles were sutured, and the skin closed with wound clips. Upon completion of the surgery, animals received a single injection of slow release buprenorphine (0.05 mg/kg, s.c.), then saline (5ml, s.c.) and Baytril (15mg/kg, s.c.) were given daily for 7 days to prevent dehydration and infection, respectively. Bladders were expressed manually at least twice daily until the voiding reflex returned.

### Exercise regimen

Beginning 4-5 days post-injury, exercised groups received 20 min of daily cycling, 5 days/week until completion of the study. No exercise was provided the day of the terminal experiment, so that the last exercise session took place >24h beforehand. Animals were secured in a support harness with the hindlimbs hanging and the feet fastened to pedals with surgical tape. The hindlimbs went through a complete range of motion during pedal rotation (45 rpm). Although the movement of the hindlimbs is passively generated by a custom-built motor-driven apparatus (Houle et al., 1999; Côté et al., 2011; Côté et al., 2014), this exercise protocol evokes rhythmic activity in both flexor and extensor muscles (Beverungen et al., 2019).

### Drug Preparation and administration

The KCC2 enhancer CLP257 (generous gift from Dr. Y. de Koninck, Université Laval, Qc, Canada) was re-suspended in dimethyl sulfoxide (DMSO) as 100mM stock solution, then was freshly diluted to 100μM in saline immediately before administration. The KCC2 inhibitor VU0240551 (Cat# 3888, Tocris Bioscience, Minneapolis, MN) was prepared from a 50mM stock solution re-suspended in DMSO and diluted to 30μM in saline immediately before administration during the terminal experiment.

### Terminal experiment

Five weeks post-injury, rats were anesthetized with isoflurane (1-4%) in O_2_ for the terminal experiment. The lumbar enlargement of the spinal cord was exposed, and the dura carefully removed. Using skin flaps of the back, an agar bath was created, and a small window was made in the solidified agar above the exposed lumbar enlargement (Gagnon et al., 2013) (**Fig. 1C**). The agar bath was filled with saline for baseline recordings and refilled with 0.5mL of the CLP257 solution for post-drug testing (or saline for control). As the half-life of CLP257 is ~15 minutes and is known to take effect at this time point (Gagnon et al., 2013), post-treatment recordings were collected 15 minutes after the application of CLP257/saline. The tibial nerve of the right hindlimb was dissected free and fitted with a cuff electrode for stimulation. The triceps surae (TS) muscles were isolated, the Achilles tendon severed distally, and tied to the lever of a servo-motor muscle puller (**Fig. 1A**) (Aurora Scientific, Aurora, ON, Canada). Bipolar wire electrodes (Cooner Wire, Chatsworth, CA) were inserted into the lateral gastrocnemius (LG) to record EMG activity in response to stretches of the TS (**Fig. 1B**), and into the interosseous muscle for H-reflex recordings (**Fig. 1D**). Muscle Force, muscle length and EMG activity obtained during the experiment were amplified (100-1000x; A-M Systems, Carlsborg, WA), band-pass filtered (10-5,000Hz) and the signal was digitized (10kHz) before being stored on a computer for analysis in Signal version 6 or Spike2 version 8 (Cambridge Electronic Design Limited, Cambridge, UK).

**Figure 1:**
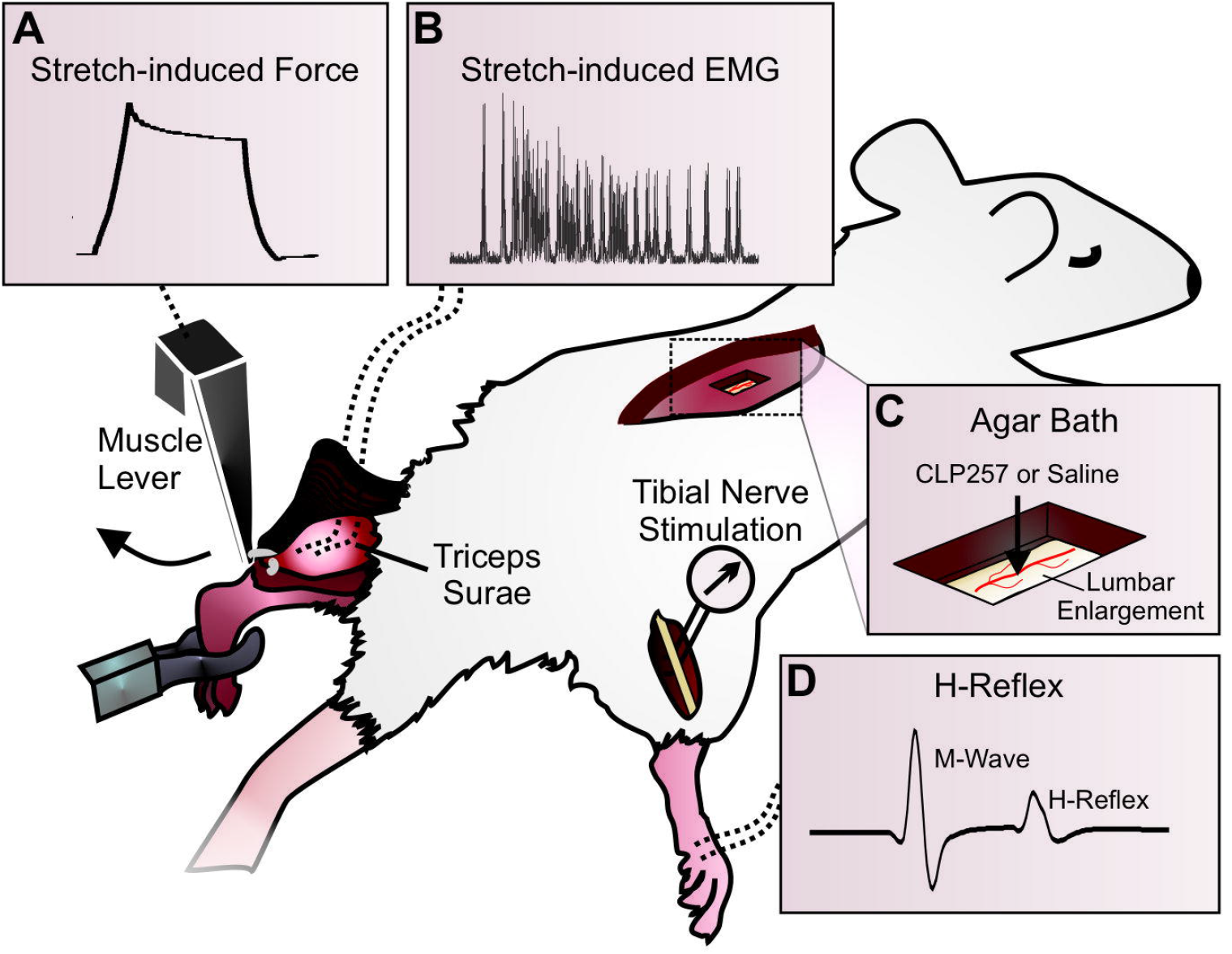
Experimental Setup. **A)** A dual mode muscle lever is attached to the left Achilles tendon to stretch the isolated triceps surae at varying lengths and speeds. The resultant force is recorded throughout the experiment. **B)** Bipolar electrodes are inserted into the lateral gastrocnemius (LG) muscle to measure EMG in response to physiological stretches. **C)** Window cut into an agar bath just above exposed lumbar enlargement for drug application. **D)** The H-reflex is evoked by a stimulation to the tibial nerve and recorded in the interosseous muscles of the right leg.

### H-Reflex

The H-reflex was evoked in the interosseus muscles by stimulating the tibial nerve with an isolated pulse stimulator delivering single bipolar pulses (100μs, A-M Systems). H-reflexes and M-waves were first recorded in response to a range of increasing stimulus intensities to determine the threshold for activation of the reflex (H-reflex threshold) and the motor response (MT). A recruitment curve was plotted by expressing the peak-to-peak amplitude of the H-reflex and M-wave responses as a function of stimulus intensity. The maximal amplitude of the H-reflex (H_max_) and the amplitude of the response of all motor units with supra-maximal stimulation of the tibial nerve axons (M_max_) were determined. Response latencies for the H-reflex and M-wave, the H_max_/M_max_ ratio, and the H_max_/ M threshold ratio were measured before and 15 minutes after treatment.

The rate-dependent depression (RDD) of the H-reflex was estimated as described previously (Côté et al., 2014; Beverungen et al., 2019). Briefly, the stimulation intensity that evoked approximately 70% of the H_max_ response was used for series of 20 consecutive stimulations at 0.3, 5 and 10Hz. The 0.3Hz series was then repeated to verify that the M-wave amplitude was still within 95% of the initial trial. The first five responses in a stimulation series were discarded to allow for reflex stabilization, and the peak-to-peak amplitude of the remaining 15 responses was averaged for each animal and frequency. The change in H-reflex response at 5Hz and 10Hz was calculated as a percentage of the response measured at 0.3Hz. The properties of the H-reflex and M-wave and the RDD were assessed before, and 15 minutes after application of CLP257 or saline onto the exposed spinal cord. All data are presented as mean ± standard error of the mean (SEM).

### Stretch-Induced Reflex and Muscle Force

The ankle joint was held at 90° flexion and the muscle was stretched approximately 1mm from its non-stretched length to achieve a baseline tension of 0.3 Newtons, which corresponds to the muscle length of the animal with an intact Achilles tendon in this joint position. Passive and reflex-mediated forces in the triceps surae was assessed by using a protocol similar to other studies in rodents (Pingel et al., 2016) and studies in humans (Lorentzen et al., 2010; Willerslev-Olsen et al., 2013). Data were collected in length servo-mode, with computer driven TS muscle stretches evoked while monitoring muscle force, muscle length, and EMG activity. In order to examine the effect of varying magnitudes of the stretch reflex, three different stretch amplitudes were applied to the TS muscle (1, 2, 3 mm) with rising phases of 5, 20 or 50ms, resulting in 9 different velocities ranging from 20 to 600mm/s. In all cases, the hold phase was 200ms, and the release phase was 100ms. Muscle force and EMG activity were recorded from the LG muscle in response to a series of ramp-hold-release stretches repeated at 4-s intervals. A second series of stretches was collected 15 minutes after application of CLP257, or saline for Controls.

Two measures of TS force output were averaged across trials and for each condition: 1) the peak force; 2) the plateau force. The peak force, which occurred at the peak of the ramp stretch, was measured as the peak-to-peak amplitude of the force evoked by a given stretch velocity. The plateau force was determined as the average force amplitude over the final 100ms of the hold phase of the stretch, before release.

EMG recordings were analyzed in response to the stretch of 600mm/s velocity (3mm amplitude, 5ms rising time) as this parameter reliably yielded the most discernable reflex responses. EMG recordings were rectified and band-pass filtered 70Hz-2500Hz offline for analysis. Muscle stretch evoked a phasic response that was measured as the integrated area of a window within the first 15ms of the stretch. An additional tonic component, defined as EMG activity that persisted >15ms after stretch onset, was also quantified when present, using the integrated area from 15ms to the end of the hold phase of the stretch.

The reflex-mediated force was also measured to confirm results of phasic EMG analysis. Approximately 40ms following the onset of the stretch, a clear increase in force is observed whenever a stretch reflex is present in the EMG (**Fig. 5A**). This additional force response, elicited by a stretch reflex contraction of the muscle, represents the reflex-mediated force (Pingel et al., 2016). Reflex-mediated forces were measured as the peak-to-peak amplitude between the peak of the reflex-mediated force and the beginning of the hold plateau. This amplitude was then normalized to the peak force amplitude elicited by that stretch. As with EMG recordings, this was solely analyzed for the stretch of 600mm/s velocity (3mm amplitude, 5ms rising time).

### Spontaneous spiking recordings and analysis

In some animals, EMG activity in LG muscle was recorded continuously and the effect of CLP257 (n=5) or saline (n=4) on spontaneous spiking activity assessed. Using a template matching function (Spike2, CED), individual motor units were discriminated based on spike shape, amplitude, and width, and confirmed with visual inspection. Spike count was averaged for each motor unit over five-minute intervals before (baseline), and up to 20 minutes after application of CLP257 or saline. In two animals (12 firing motor units), CLP257 was then washed off the exposed lumbar enlargement 20 minutes following its application and was replaced with the KCC2 inhibitor VU0240551. The average spike count following application of VU0240551 was then measured to determine if the effects of CLP257 on spiking could be attributed to changes in KCC2 function. In addition to spiking frequency, the number of recruited motor units was also estimated. A motor unit was termed as “recruited” when the template matching software in Spike2 plotted a unit exclusively after application of terminal treatment. Care was taken to validate that each recruited unit was in fact separate from those that fired during baseline recordings.

### Immunofluorescent staining

Rats were sacrificed using an overdose of Euthasol (390 mg/kg sodium pentobarbital and 50mg/kg phenytoin, i.p.) immediately following the terminal experiment, and perfused transcardially with cold saline followed by 4% paraformaldehyde in PBS. The lumbar spinal cord (L1–L6) was harvested, post-fixed overnight, and transferred to 30% sucrose in PBS for cryoprotection. Spinal cord tissue was then sectioned transversally (25μm) on a cryostat and mounted on slides for immunostaining. Sections were first pre-incubated in BSA (1%), donkey serum (10%) and 0.2% Triton x-100 OX in PBS for one hour at room temperature. Sections were then incubated with the following primary antibodies at 4°C overnight: rabbit anti-KCC2 (1:1000, Cat#07-432, Millipore, RRID: AB_310611) and goat anti-ChAT (1:100, Cat#AB144P, Millipore, RRID: AB_2079751). Species-specific secondary antibodies (1:1000, goat anti-rabbit conjugated to Alexafluor 488 for KCC2, and donkey anti-goat conjugated to rhodamine for ChAT; Jackson ImmunoResearch) were applied for 2h at room temperature, and the slides were cover-slipped before imaging. Spinal cord sections were imaged using a Leica DM550B fluorescent microscope and a Retiga-SRV digital color camera (QImaging) controlled by Slidebook imaging software version 6.0.11 (Olympus). Eighteen lumbar motoneurons were averaged per group (6 motoneurons for each animal, n=3 per group), and were identified with ChAT+ labeling, typical large size, and location within the ventral horn. The fluorescence intensity of KCC2 immunolabelling for the motoneuronal membrane was measured using ImageJ software and averaging the integrated area of the density curve obtained by drawing three lines across each motoneuron (yielding six data points per cell) (Boulenguez et al., 2010; Côté et al., 2014; Liabeuf et al., 2017). To examine the amount of KCC2 expressed in the membrane compared to the cytosol, mean pixel intensities for membranes were normalized to intensities in the cytoplasm for each motoneuron (Boulenguez et al., 2010).

### Statistical Analysis

Paired t-tests were used to compare differences in H and M maximum amplitudes, H and M latencies, H and M thresholds, the H_max_/M_max_ ratio, and the H_max_/M threshold. These were compared for each animal before and after receiving either CLP257 or Saline. Where data was not normally distributed, a Wilcoxon test was used. For RDD measurements, three-way mixed ANOVAs followed by Holm-Sidak *post-hoc* tests were used to determine significant differences in H-reflex amplitude across stimulation frequencies (5Hz and 10Hz, normalized to 0.3Hz; within-subject factor), treatments (CLP257/Saline; between subject factor) and condition (pre and post treatment; within-subject factor). This was done separately for un-exercised and exercised groups. To compare effects of CLP257 on the RDD in un-exercised animals to effects of exercise, a two-way mixed ANOVA was run with a Holm-Sidak *post-hoc* test. Differences in H-reflex amplitude were compared across stimulation frequencies (5Hz and 10Hz, normalized to 0.3Hz; within-subject factor) and groups (un-exercised + CLP257 or exercised; between-subjects factor).

A two-way mixed ANOVA was used to compare both peak forces and plateau forces between un-exercised and exercised groups (between-subjects factor) for 9 different stretches of the triceps surae (within-subjects factor). Two-way ANOVAs were also used to determine the effects of CLP257 (pre and post CLP257; within-subjects factor) on these force outputs for the different stretches (within-subjects factor).

Phasic and tonic EMG responses following treatment were averaged across 10 trials and normalized to the pre-treatment response (averaged over 10 pre-treatment trials) in each animal. Unpaired t-tests were then used to compare normalized post-treatment responses between CLP257 and saline treated animals. This was done separately for un-exercised and exercised groups.

To confirm results of phasic EMG responses, reflex-mediated forces were measured and normalized to the peak force elicited by that stretch. Ten reflex-mediated forces were measured and averaged both before and after treatment in each animal. Using a two-way mixed ANOVA, averages were then compared across treatments (CLP257/Saline; between-subject factor) and condition (pre and post treatment; within-subject factor).

One-way ANOVAs were used to compare mean pixel intensities of immuno-labeled KCC2 between groups. Unless otherwise stated, values were normally distributed as assessed by Shapiro-Wilk’s test (*p* > 0.05) and Normal Q-Q plot, and there were no outliers in the data, as assessed by inspection of a boxplot and by examination of studentized residuals for values greater than ±3. There was homogeneity of variances, as assessed by Levene’s test of homogeneity of variance (*p* > 0.05). Unless stated otherwise, Mauchly’s test of sphericity indicated that the assumption of sphericity was met (*p* > 0.05). Where it was violated, a Greenhouse-Geisser correction was applied. All data are reported as mean ± SEM. Significance level was set to *p* < 0.05. Significance of a simple two-way interaction and a simple-simple main effect was accepted at a Bonferroni-adjusted alpha level of 0.025.

To analyze the effect of CLP257 on spike count, we used a generalized linear mixed model with a negative binomial distribution because the data was not normally distributed. Fixed effects for time, treatment and their interaction were included. Using a stacked random effect for each motor unit and animal, the model controlled for inter-individual differences. The glmmTMB package (Brooks et al., 2017) for R 3.2 was used to fit the model and account for zero-inflation and overdispersion. Backwards elimination was used starting with the full factorial model for the random effect, zero inflation and dispersion terms. Effects were removed from the model if the likelihood-ratio test was not significant. The intercepts for the random effect and the zero-inflation remained in the model, along with the dispersion terms (time and treatment). Q-Q plots and histograms of Pearson’s residuals were used to confirm negative binomial distribution of the residuals. Post-hoc comparisons using the Tukey’s test were used to assess spike count following VU0240551 application compared to CLP257 application.

## Results

We have previously shown that restoring KCC2 activity after SCI is critically involved in preventing the development of spasticity following activity-based therapies (Côté et al., 2014; Beverungen et al., 2019). Given that many patients present a significant number of co-morbidities that render them unable to participate in a rehabilitation program, particularly in the acute phase (Simon and Yelnik, 2010), our objective was to investigate if pharmacologically restoring chloride homeostasis can be used as an alternative to reduce symptoms of spasticity after chronic SCI.

### H-reflex modulation is improved by restoring chloride homeostasis after chronic SCI

We first investigated the effect of restoring chloride homeostasis on hyperreflexia and spasticity after chronic SCI by measuring H-reflex excitability before and after application of the KCC2 enhancer, CLP257, to the exposed spinal lumbar enlargement five weeks post-injury (**Fig. 1C**). Following stimulation to the tibial nerve, two responses can be recorded in the interosseous muscle: 1) the M-wave, which is the response to the direct activation of motor axons, and 2) the H-reflex, which results from the activation of Ia afferents that directly contact motoneurons (**Fig. 1D**). Table 1 displays the general properties of the M-wave and H-reflex such as maximal amplitude, latency and threshold for activation before and 15 minutes after CLP257 or saline application. Paired t-tests found that CLP257 did not significantly alter H or M latencies or thresholds after chronic SCI (p > 0.05). To assess the proportion of motoneurons recruited through the mono-synaptic reflex compared to the activation of the whole motor pool, the H_max_/M_max_ ratio was measured. This did not change significantly after application of either CLP257. Similarly, the H_max_/M threshold ratio, an estimate of the relative activation of the motor pool required to reach maximal reflex amplitude, also showed no significant difference after saline or CLP257 application (**Table 1)**.

We then measured the effect of restoring chloride homeostasis after chronic SCI on the rate-dependent depression (RDD) of the H-reflex. In the intact spinal cord, the amplitude of the H-reflex is decreased by the repetitive activation of Ia afferents (Crone and Nielsen, 1989; Thompson et al., 1992; Hultborn et al., 1996). SCI disrupts reflex modulation and leads to a decrease in the RDD in animals (Thompson et al., 1992) and in humans (Calancie et al., 1993; Trimble et al., 2001; Grey et al., 2008). Impaired RDD after chronic SCI has been associated with decreased KCC2 activity and subsequent depolarization in E_Cl−_ that contributes to hyperreflexia and spasticity (Boulenguez et al., 2010; Bos et al., 2013). To assess if CLP257 improves RDD after chronic SCI, H-reflexes were evoked at 0.3Hz, 5Hz and 10Hz before and after application. Figure 2A shows representative traces of the H-reflex evoked by a 10Hz stimulation train in the same animal before and 15 minutes after the application of CLP257. In this animal, CLP257 markedly decreased the amplitude of the H-reflex as compared to baseline. A three-way mixed ANOVA revealed a statistically significant interaction between treatment (CLP257/saline), and condition (pre/post treatment) (F_(1,16)_ = 11.11, *p* = 0.004). There was a significant difference in H-reflex amplitude across stimulation frequencies (F_(1,16)_ = 23.32, *p* < 0.001), suggesting that some level of depression is still present after chronic SCI whether CLP257 was applied or not. However, within-subjects contrasts found a significant effect of condition (pre/post treatment) on H-reflex depression in CLP257 treated animals (n=10) (F_(1,9)_ = 31.72, *p* < 0.001) but not in saline treated animals (n=8) (F_(1,7)_ = 0.04, *p* = 0.839) (**Fig. 2B**), suggesting that CLP257 affected the depth of depression.

**Figure 2:**
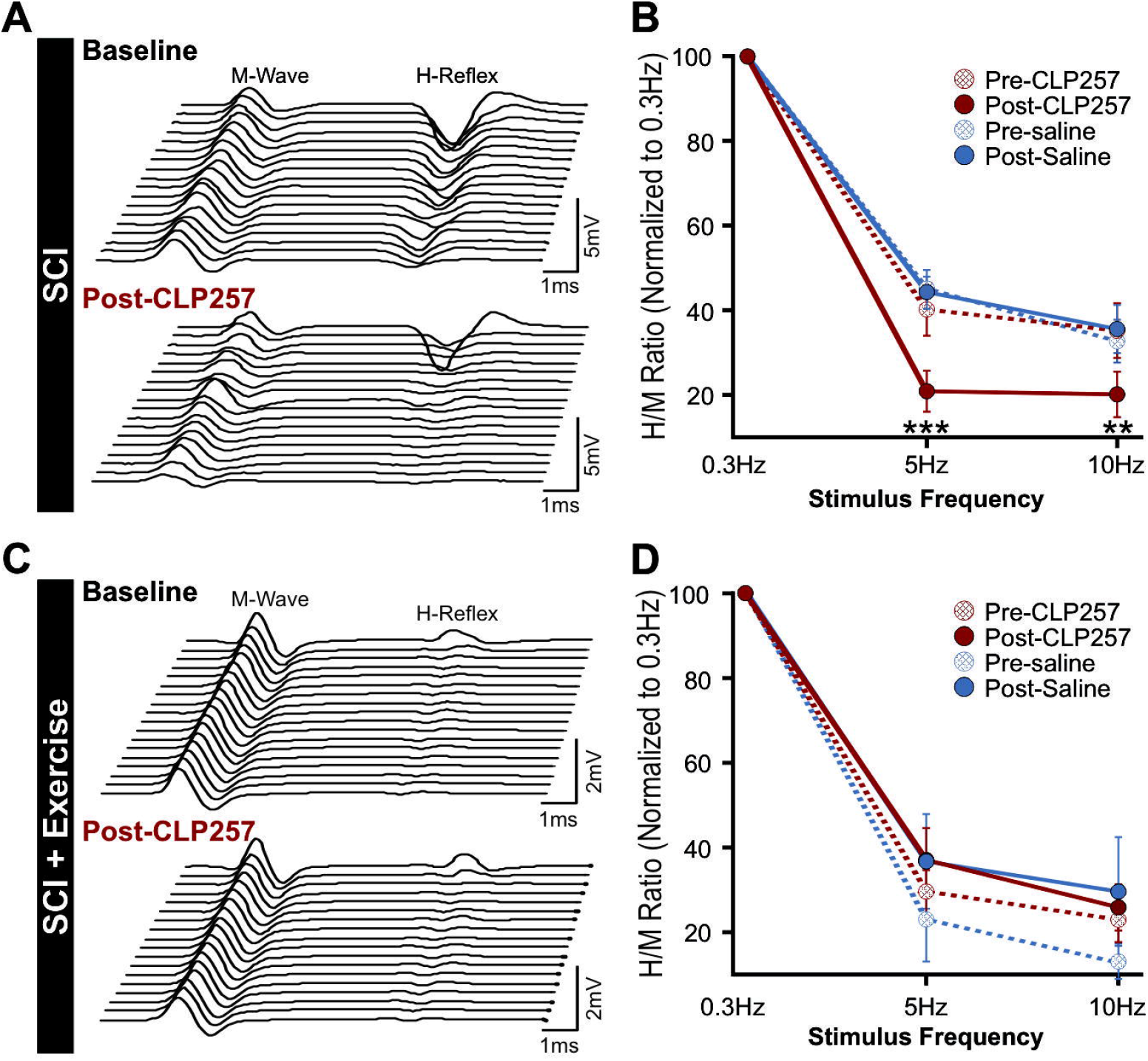
Enhancing KCC2 activity restores the frequency-dependent depression of the H-reflex after chronic SCI. **A)** Representative H-reflex recordings over a series of 20 stimulations to the tibial nerve at 10Hz from a chronic SCI animal before (top) and after CLP257 administration (bottom). CLP257 dramatically decreased H-reflex amplitude and increased the rate-dependent depression of the H-reflex. **B)** There was a statistically significant interaction between treatment (CLP257/saline) and condition (pre/post treatment) in chronic SCI animals (F_(1,16)_ = 11.11, *p* = 0.004). CLP257 decreased H-reflex amplitude at 5Hz (t_(9)_ = 6.50, *p* < 0.001) and 10Hz (t_(9)_ = 3.91, *p =* 0.004), whereas saline had no effect at either 5Hz (t_(7)_ = 0.20, *p* = 0.844) or 10Hz (t_(7)_ = −0.50, *p* = 0.633). **C)** Representative H-reflex recordings over a series of 20 stimulations to the tibial nerve at 10Hz from a chronic SCI animal that was exercised for 4 weeks, before (top) and after CLP257 administration (bottom). CLP257 showed no noticeable effect on the H-reflex modulation. **D)** Overall, there was no significant interaction between frequency, treatment, and condition (*F*_(1, 14)_ = 2.273, *p* = 0.154), nor was there a significant interaction between condition and treatment (*F*_(1, 14)_ = 1.251, *p* = 0.282). **p<0.01 ***p<0.001.

Overall, CLP257 decreased the H-reflex amplitude from 42 ± 6% to 21 ± 5% at 5Hz (t_(9)_ = 6.50, *p* < 0.001) and from 36 ± 6% to 21 ± 5% at 10Hz (t_(9)_ = 3.91, *p =* 0.004) as compared to 0.3Hz. Saline however, had no effect, with values similar to baseline at 5Hz (t_(7)_ = 0.20, *p* = 0.844) and 10Hz (t_(7)_ = −0.50, *p* = 0.633). These results suggest that CLP257 improved reflex modulation and improved the RDD after chronic SCI.

### Pharmacologically restoring chloride homeostasis parallels the effect of exercise on the H-reflex after chronic SCI

We have previously shown that the impairment in RDD is prevented by activity-based therapies (Côté et al., 2014) via a BDNF-dependent increase in KCC2 expression on lumbar motoneuronal membranes (Beverungen et al., 2019). To determine if CLP257 further ameliorates reflex excitability and modulation in animals undergoing a rehabilitation program, a group of rats received bike-training for 4 weeks following SCI. The general features of the M-wave and H-reflex before and 15 minutes after CLP257 or saline application are displayed in Table 1. CLP257 did not significantly alter any of these parameters (p > 0.05), with both M-wave and H-reflex properties similar post-CLP257/saline as compared to their respective baseline values. We then measured the effect of restoring chloride homeostasis on H-reflex amplitude at increasing stimulus frequencies in exercised animals. To assess if CLP257 improves RDD in exercised animals following chronic SCI, H-reflexes were evoked at 0.3Hz, 5Hz and 10Hz before and after treatment application. Figure 2C shows representative traces of the M-wave and H-reflex in the same animal in response to a 10Hz stimulation train before and 15 minutes after application of CLP257 to the lumbar enlargement. CLP257 had no further effect on the amplitude of the H-reflex in this animal, as a decrease in amplitude was already present before drug application (Côté et al., 2014; Beverungen et al., 2019). A three-way mixed ANOVA was run to compare the effects of CLP257 (n=11) or saline (n=5) on H-reflex amplitude at increasing stimulation frequencies. There was a statistically significant main effect of stimulation frequencies (F_(1,14)_ = 13.48, *p* = 0.003). However, there was no statistically significant interaction between stimulation frequency, treatment (CLP257/saline), and condition (pre/post treatment) (F_(1,14)_ = 2.27, *p* = 0.154). There was also no significant two-way interaction between condition and treatment (F_(1,14)_ = 1.25, *p* = 0.282). Overall, CLP257 did not significantly affect the amplitude of the H-reflex in exercised animals (**Fig. 2D**), with similar values pre- and post-CLP257 at 5Hz (31 ± 5 % vs. 38 ± 7%) (t_(10)_ = −1.58, *p* = 0.146) and 10Hz (25 ± 5% vs. 27 ± 5%) (t_(10)_ = −0.54, *p* = 0.602).

We further investigated if exercise and CLP257 have similar effects in restoring the RDD after chronic SCI. A two-way mixed ANOVA found no significant main effect of treatment (CLP257 vs. Exercise) on H-reflex depression (F_(1,20)_ = 1.02, *p* = 0.326). This indicates there was no significant difference in H-reflex depression in chronic SCI animals treated with CLP257 compared to those that had followed an exercise program. The amplitude of the H-reflex was similar between SCI + CLP257 and SCI + Ex at 5Hz (22 ± 5% vs. 31 ± 5 %) (t_(40)_ = 1.36, *p* = 0.328) and at 10Hz (21 ± 5% vs. 25 ± 5%) (t_(40)_ = 0.62, *p* = 0.539). This suggests that pharmacologically increasing KCC2 activity can reduce hyperreflexia to similar levels observed following a rehabilitation program.

### Restoring chloride homeostasis does not affect stretch-induced muscle force after SCI

To further assess how increasing KCC2 activity decreases hyperreflexia, we evaluated the force and EMG output in response to stretches of the triceps surae (TS) at 9 different velocities. Figure 3 shows representative force traces in response to stretches of the muscle. For slow stretches with 50ms rising time, the force increases linearly with the stretch until reaching a small peak of maximal force, then plateauing during the hold phase of the stretch (**Fig. 3A**). This peak is more pronounced with stretches of 20ms rising time, and even more so with stretches of 5ms rising time (**Fig. 3B-C**). For each stretch, the amplitude of the peak force response and the average amplitude of the plateau force was calculated. There was no significant difference in stretch-induced forces in exercised and un-exercised animals, for both peak and plateau forces (F_(1,6)_ = 3.33, *p* =0.118, and F_(1,4)_ = 0.04, *p* = 0.845, respectively, two-way ANOVA, data not shown); therefore, exercised and un-exercised animals were pooled together for further analysis.

**Figure 3:**
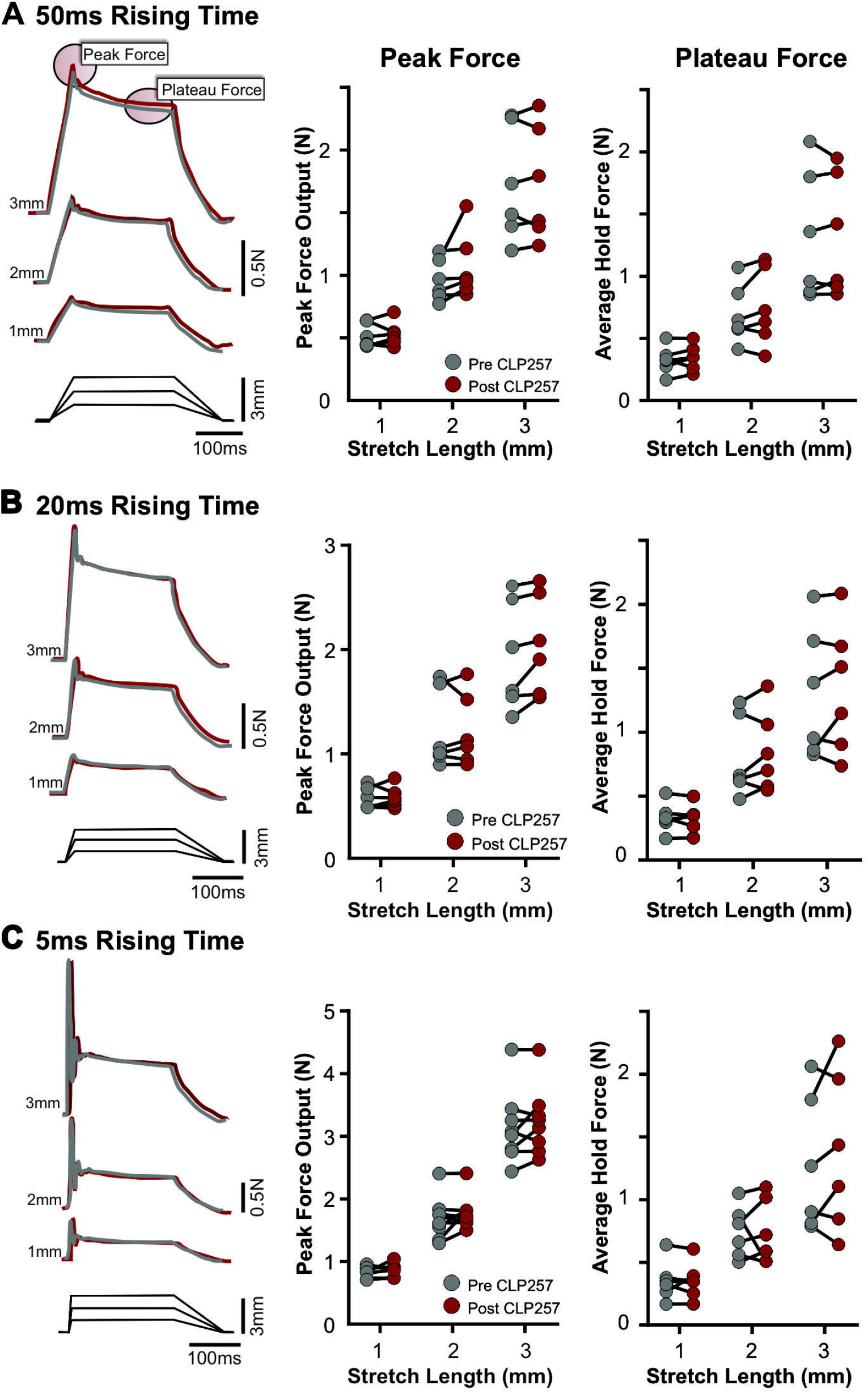
CLP257 does not affect stretch-induced force responses after SCI. Forces exerted in response to 1, 2, or 3mm stretches of the triceps surae with rising phases of 50ms (**A**), 20ms (**B**) and 5ms (**C**). Representative traces are from a single animal before (grey) and after (red) application of CLP257. Shown in the insets are the peak force and the plateau force (the mean force during the last 100ms of the hold phase). Each graphed data point represents the average peak force (left) and plateau force (right) induced in each animal for each stretch averaged over 10 trials, before and after application of CLP257. With each stretch, CLP257 application did not significantly affect either the peak force (F_(8,45)_ = 3.92, *p* = 0.054), or the plateau force (F_(1,45)_ = 2.722, *p* = 0.1059).

Two-way ANOVAs found no statistically significant main effect of condition (pre/post treatment) on peak force values (F_(1,45)_ = 3.92, *p* = 0.054). There was also no significant main effect of condition on the average plateau forces (F_(1,45)_ = 2.72, *p* = 0.106). This indicates there was no effect of CLP257 on peak and plateau forces with either 1mm, 2mm, or 3mm stretches with rising times of 50ms (**Fig. 3A**), 20ms (**Fig. 3B**) or 5ms (**Fig. 3C**).

### CLP257 reduces the phasic and tonic stretch-induced EMG responses after SCI

While the H-reflex was assessed in the right hindlimb via tibial nerve stimulation, in the left hindlimb we assessed reflexes in a more holistic manner via mechanical stretching of the triceps surae (TS) muscles. After chronic SCI, quick muscle stretches elicit a large phasic EMG response, with a latency corresponding to a monosynaptic reflex. An exaggerated monosynaptic reflex is characteristic of SCI and is thought to contribute to the production of clonus and hyperreflexia (Nielsen et al., 2007). In addition to exaggerated phasic responses, long-lasting responses can also be present (Bennett et al., 1999; Bennett et al., 2004; Li et al., 2004; Murray et al., 2011b) and contribute to triggering spasms in SCI patients (Norton et al., 2008).

We evaluated the effect of restoring chloride homeostasis with CLP257 on LG muscle EMG activity evoked by TS muscle stretches. When slow stretches were applied, little or no EMG activity was elicited (*not shown*). However, faster stretches with a velocity of 600mm/s (3mm, 5ms rising time) evoked large phasic responses (**Fig. 4A**) with an average latency of 4.35 ± 0.1ms. In addition to exaggerated phasic responses, a considerable number of chronic SCI animals (n= 7/17) displayed robust and long-lasting EMG activity with an onset 1.5-15ms after the phasic response had terminated, and in some cases persisted for up to 3.5 seconds after the muscle stretch ended, suggesting the contribution of a polysynaptic pathway. Figure 4A shows representative traces of the phasic and tonic EMG responses in a chronic SCI animal before and after application of CLP257. The amplitude of the phasic response was reduced by CLP257 (n=4), but not by saline (n=6) after chronic SCI. An unpaired t-test found that phasic responses were significantly lower after CLP257 treatment (55.6 ± 13.0% of baseline amplitude) compared to saline treatment (99.4 ± 7.1% of baseline amplitude) (t_(8)_ = 3.24, *p* = 0.012) (**Fig. 4B**). Tonic responses, when present, were also significantly reduced by CLP257 treatment compared to saline treatment (t_(5)_ = 3.337, *p* = 0.0206). CLP257 reduced responses to 46.6% ± 18.3% of baseline values, while responses in saline treated animals remained at 101.2 ± 4.9% of baseline values (**Fig. 4B**). This demonstrates that increasing KCC2 activity robustly attenuates long polysynaptic reflexes that initiate muscle spasms. Figure 4C shows representative EMG responses to stretch in exercised animals after a chronic SCI. Interestingly, tonic EMG responses were not observed in any animals that had received bike-training (n=13) (**Fig. 4C**). Contrary to un-exercised animals, CLP257 had no significant effect on the amplitude of the phasic component of the stretch reflex (t_(5)_ = 0.063, *p* = 0.9523) (**Fig. 4D**). This again suggests that CLP257 parallels the effects of exercise on hyperreflexia.

**Figure 4:**
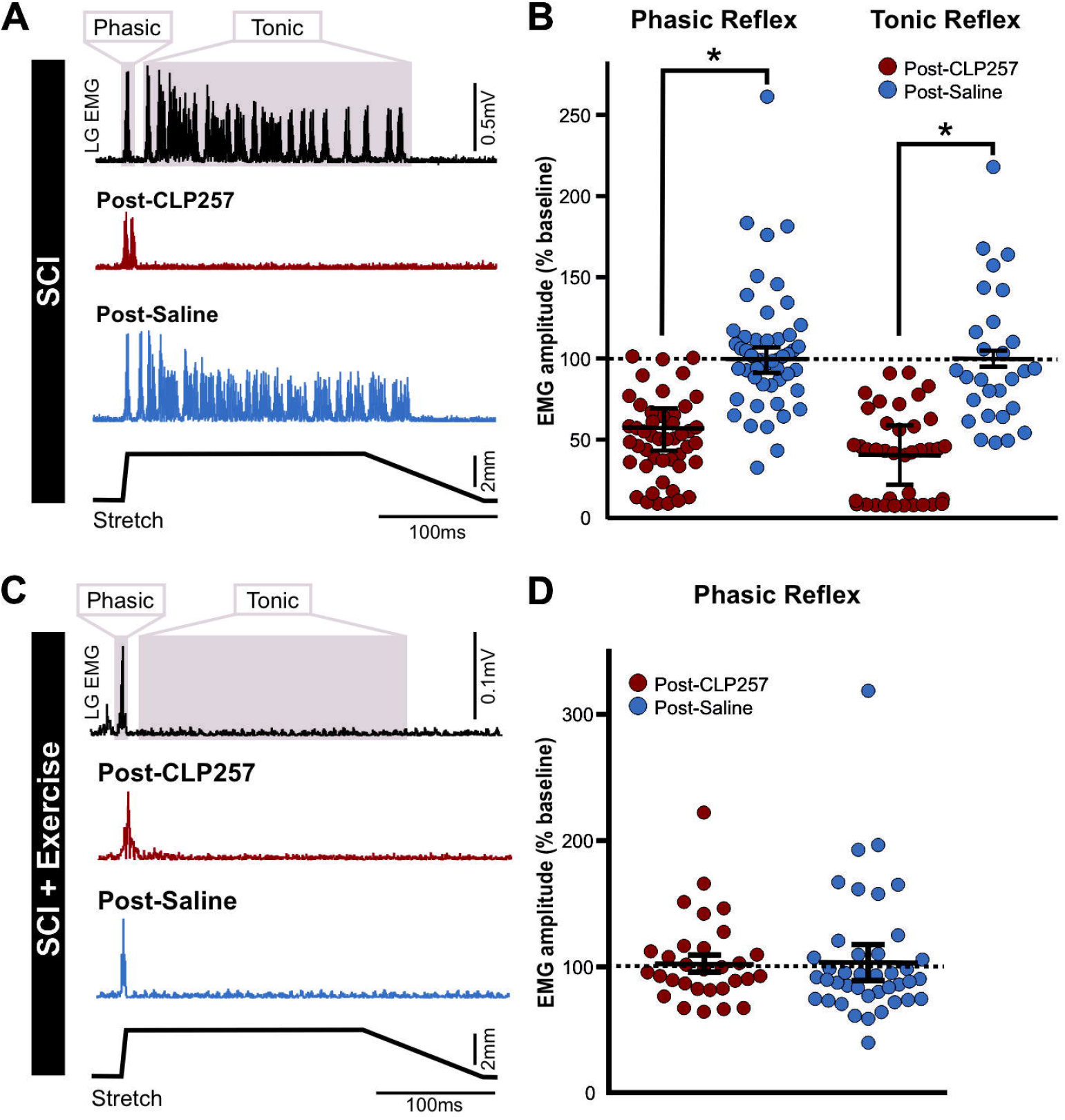
Enhancing KCC2 activity reduces stretch-induced EMG responses after chronic SCI. After chronic SCI, EMG responses to stretch have a phasic (early, short-lasting) and/or tonic (late, long-lasting) component. **A)** CLP257 decreases the amplitude of the phasic response after chronic SCI and prevents the development of a tonic response. **B)** Phasic responses following CLP257 treatment were significantly reduced compared to saline treatment (t_(8)_ = 3.238, *p* = 0.012) in chronic SCI animals. Similarly, the tonic response was also significantly reduced by CLP257 compared to saline treatment (t_(5)_ = 3.337, *p* = 0.0206). **C)** Exercise prevented the development of tonic EMG activity in LG muscle in response to triceps surae muscle stretch, and CLP257 did not alter the phasic response. **D)** CLP257 had no significant effect on the amplitude of the phasic response in exercised animals, with values similar to saline values (t_(5)_ = 0.063, *p* = 0.952). *p<0.05.

### CLP257 reduces the reflex-mediated force

In addition to measuring EMG responses, we evaluated the reflex-mediated force to ensure that differences in EMG read-outs observed with CLP257 were not due to movement of the wire electrodes. The reflex-mediated force response is seen as a clear increase in force, at an approximate latency of 40ms, elicited by a stretch reflex contraction of the muscle (**Fig. 5A**)(Pingel et al., 2016). The delay of this increase in force compared with the latency of the EMG response is explained by the electromechanical delay. In agreement with our stretch-induced EMG results, there was a significant interaction between treatment (CLP257/saline) and condition (pre/post treatment) (F_(1,8)_ = 14.61, *p* = 0.0051, two-way mixed ANOVA). CLP257 reduced the amplitude of the reflex-mediated force to 85.4 ± 3.7 % of pre-CLP257 values (t_(8)_ = 5.70, *p* < 0.001), while saline did not affect it (t_(8)_ = 0.296, *p* = 0.775) with average forces post-saline remaining at 98.3 ± 3.8 % of pre-saline values (**Fig. 5B**).

**Figure 5:**
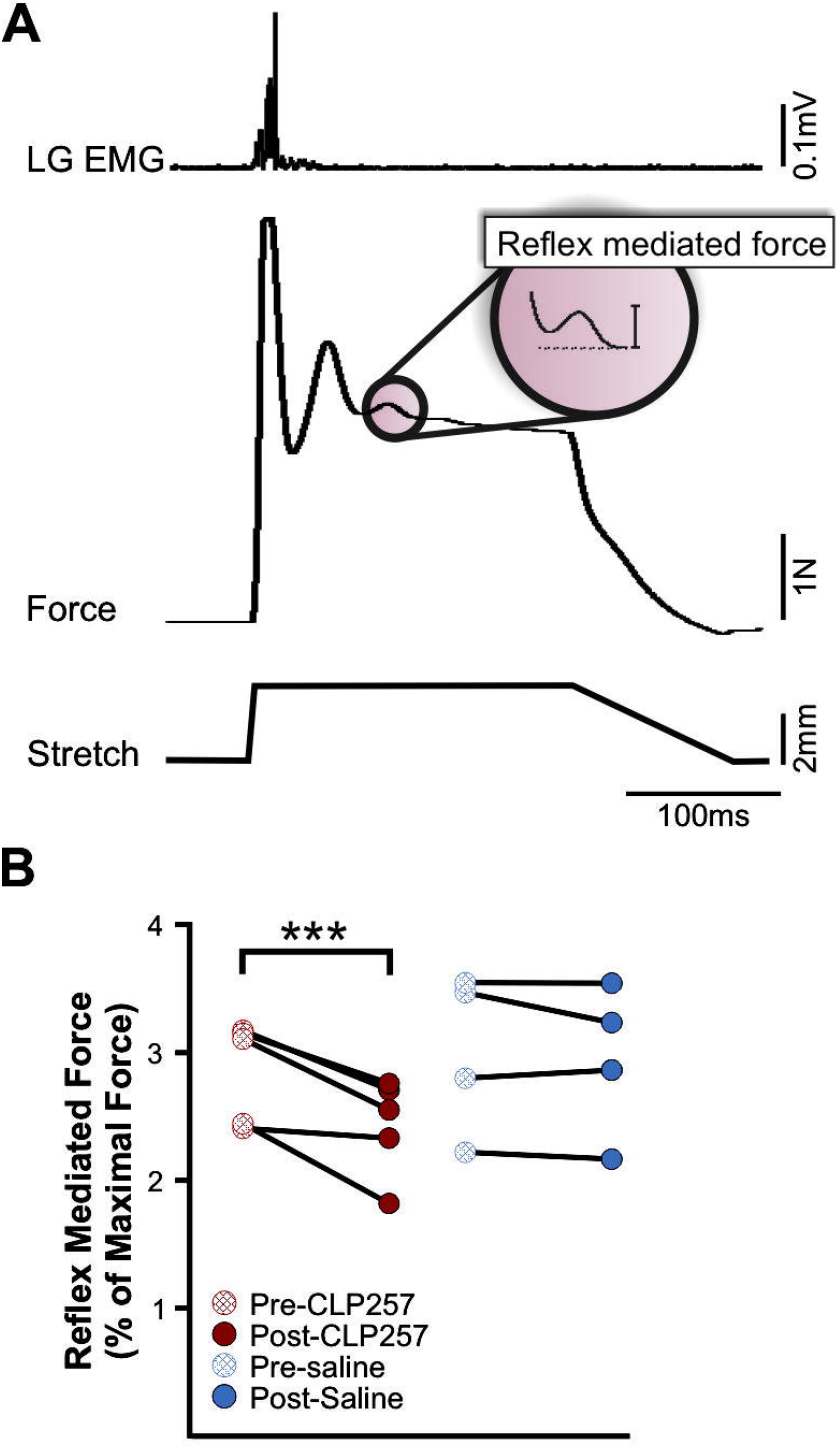
CLP257 reduces reflex-mediated forces. **A**) Representative traces of force output of triceps surae and EMG activity in LG muscle in response to a triceps surae stretch (3mm length, 5ms rising phase). Within the inset is the reflex-mediated force. **B**) There was a significant interaction between treatment (CLP257/saline) and condition (pre/post treatment) after chronic SCI (F_(1, 8)_ = 14.61, *p* = 0.0051), with CLP257 significantly reducing the amplitude of the reflex-mediated force (t_(8)_ = 5.702, *p* < 0.001), and saline having no effect (t_(8)_ = 0.2959, *p =* 0.7749). ***p<0.001.

### Exercise increases spontaneous motor-unit activity in hindlimb muscles, which is further increased by CLP257 treatment

We also monitored spontaneous motor unit spiking before and after application of either CLP257 or saline after chronic SCI. Interestingly, exercise increased the occurrence of animals displaying spontaneous spiking activity in LG (n=10/13) as compared to chronic SCI animals that did not follow a rehabilitation program (n=3/17) (χ^2^_(1, N = 30)_ = 8.265, *p* = 0.004, odds ratio = 4.36) (**Fig.6A**). Figure 6B shows a representative recording from an exercised rat that displayed spontaneous spiking at baseline. CLP257 increased the spiking frequency, while the KCC2 inhibitor VU0240551 greatly reduced spiking towards baseline levels **(Fig.6C)**. CLP257 had no effect on animals that did not display spontaneous spiking activity, with the EMG remaining silent after CLP257 application, whether animals were exercised (n=3) or not (n=8) (*data not shown*). Animals not displaying spontaneous activity were excluded from further analysis. A generalized linear mixed model found a significant interaction between treatment (CLP257/saline) and condition (pre/post treatment) on the average spike count per motor unit (χ^2^_(4)_ = 21.387, *p* = 0.0003). CLP257 increased average spike count at 5 minutes (t_(217)_ = 3.254, *p=* 0.011), 10 minutes (t_(217)_ = 2.895, *p =* 0.0337), 15 minutes (t_(217)_ = 3.519, *p =* 0.0048) and 20 minutes (t_(217)_ = 4.250, *p =* 0.0003) post-application. Saline treatment had no significant effect on average spiking count at any time point (*p* > 0.05), suggesting that our preparation was stable, and that time did not influence spontaneous spiking activity. There was also a significant difference in spike count between CLP257 and saline treated animals at 10 minutes (t_(217)_ = 2.338, *p =* 0.0203), 15 minutes (t_(217)_ = 2.744, *p =* 0.0066), and 20 minutes (t_(217)_ = 2.890, *p =* 0.0042) post-treatment (**Fig. 6D**). To confirm that the observed effects can be attributed to an increase in KCC2 activity, we applied VU0240551 on the lumbar spinal cord 20 minutes after receiving CLP257 treatment in a subgroup of animals (n=2). Post-hoc comparisons using the Tukey’s test determined that blocking KCC2 with VU0240551 significantly reduced mean spike count from 96.5 counts per 5 min to 12.9 counts per 5 min (t_(172)_ = 5.226, *p* < 0.0001). In fact, the mean spike count after VU0240551 application was significantly smaller than at any time point after CLP257 application (*p* < 0.005) and was not statistically significant from pre-CLP257 values (t_(172)_ = −1.480, *p* = 0.6775) (**Fig.6D**). In addition to increasing spontaneous spiking of motor units that were already active, application of CLP257 appeared to induce the recruitment of new motor units. After CLP257 treatment, four out of five animals recruited more motor units, whereas only one out of four animals recruited another unit following saline treatment (**Fig. 6E**).

**Figure 6:**
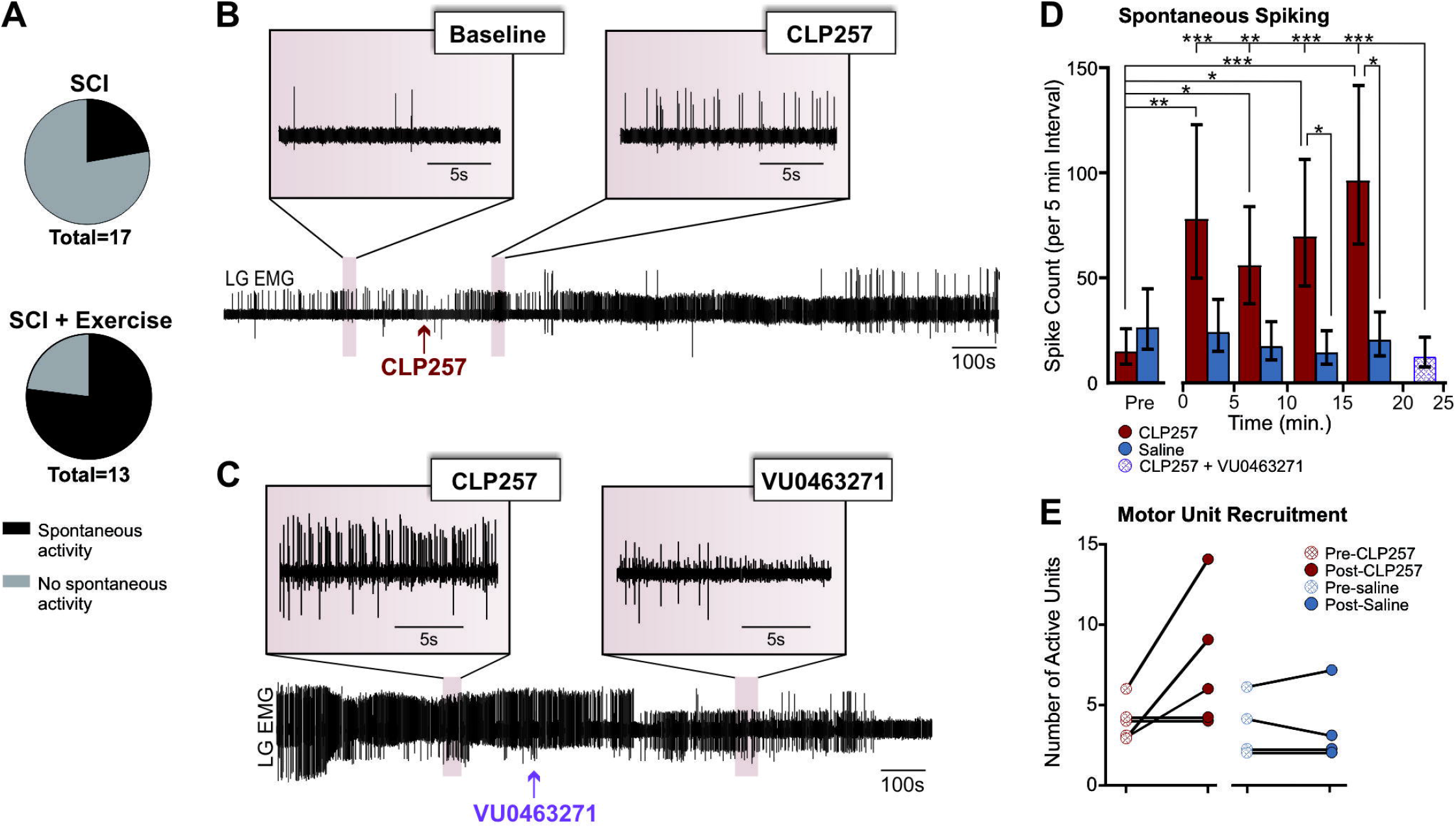
CLP257 increases spontaneous spiking activity in LG muscle after SCI. **A)** Exercise increased the occurrence of spontaneous spiking activity in the lateral gastrocnemius muscle as compared to un-exercised SCI animals (χ^2^_(1, N = 30)_ = 8.265, *p* = 0.004). **B-C)** Representative EMG recording of an animal displaying spontaneous activity at rest in LG muscle. CLP257 increased spiking frequency (**B**) while VU0240551 returned spiking activity toward baseline levels (**C**). **D)** Application of CLP257 increased spontaneous spiking activity in exercised animals (n=5) as early as 5 minutes after drug application. Blocking KCC2 with VU0240551 post-CLP257 reversed this effect (n=2). **E)** CLP257 also increased the number of motor units recruited as compared to saline application. *p<0.05. **p<0.01. ***p<0.001

### CLP257 restores KCC2 membrane expression in lumbar motoneurons after chronic SCI

KCC2 must be expressed in the neuronal membrane to extrude chloride from the cell. Following chronic SCI, there is a ~10-20% decrease in motoneuronal KCC2 expression, and the translocation of KCC2 to the somatic membrane of motoneurons is significantly reduced (Boulenguez et al., 2010; Bos et al., 2013). As CLP257 works post-translationally to increase KCC2 membrane expression (Gagnon et al., 2013; Ferrini et al., 2017), we compared the ratio of membrane KCC2 to cytosolic KCC2 immuno-labelling between groups. Figure 7 shows KCC2 immunoreactivity in lumbar motoneurons after chronic SCI in animals that received CLP257 (**Fig. 7A**) or saline (**Fig.7B**), and exercised animals that received CLP257 (**Fig.7C**) or saline (**Fig. 7D**). A one-way ANOVA found the interaction between the membrane-cytosol ratio and groups was statistically significant (F_(3,68)_ = 21.43, *p* < 0.0001). In chronic SCI rats, the ratio of KCC2 in the membrane to cytosol was significantly larger with CLP257 treatment than with saline treatment (t_(68)_ = 10.66, *p* < 0.001). CLP257 also induced a significant difference in the membrane-cytosol ratio in exercised animals (t_(68)_ = 3.84, *p* = 0.0409), and CLP257 combined with exercise significantly increased the ratio compared to saline-treated, unexercised animals (t_(68)_ = 6.98, *p* < 0.0001). Additionally, un-exercised CLP257-treated animals had a significantly higher ratio than exercised saline-treated animals (t_(68)_ = 7.52, *p* < 0.001). This suggests, as expected, that CLP257 works post-translationally by moving whatever KCC2 is left after injury from the cytoplasm to the membrane, whereas exercise increases overall KCC2 expression as well as membrane expression (Côté et al., 2014).

**Figure 7:**
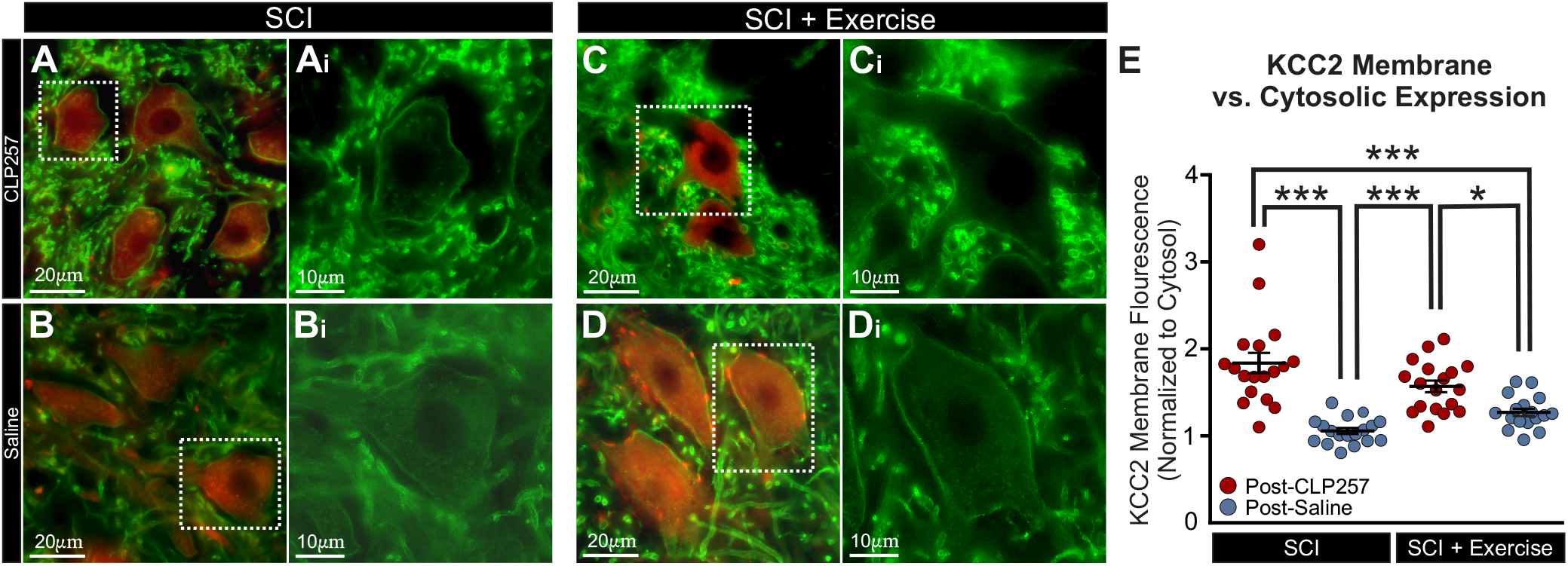
CLP257 increases KCC2 membrane expression in lumbar motoneurons. **A-D)** Representative images showing KCC2 (green) and ChAT (red) immunoreactivity in lumbar motoneurons of chronic SCI rats treated with CLP257 (A) or saline (B), and exercised rats treated with CLP257 (C) or saline (D). **E)** Mean pixel intensities of the immunolabeling were measured under 3 traces overlaying each motoneuron and normalized to a delimited area of the cytosol. There was a significant interaction between the membrane-cytosol KCC2 ratio and groups (F_(3, 68)_ = 22.17, *p* < 0.001). In both un-exercised and exercised animals, CLP257 significantly increased the amount of KCC2 expressed in the membrane vs. in the cytosol. *p<0.05, ***p<0.001.

## Discussion

It is widely accepted that multiple factors contribute to the emergence and maintenance of spastic symptoms. In particular, the contribution of increased motoneuronal intrinsic excitability and loss of descending inhibition have been thoroughly studied (Bennett et al., 2001; Nielsen et al., 2007). More recently, a disruption in chloride homeostasis was identified as another significant contributor. Under normal physiological conditions, intracellular concentration of chloride is maintained low by the chloride extruder KCC2, which favors chloride influx through GABA_A_R and membrane hyperpolarization (Kaila et al., 2014). After chronic SCI, there is a decrease in KCC2 expression in lumbar motoneurons (Boulenguez et al., 2010), likely via increased calpain-mediated proteolysis (Plantier et al., 2019). This leads to a reversal of GABA-mediated responses from hyperpolarization to depolarization, and increases the chance that a motoneuron will fire an action potential (Vinay and Jean-Xavier, 2008; Boulenguez et al., 2010). This shift in chloride homeostasis contributes to the development of spasticity, as measured by a decrease in H-reflex RDD (Vinay and Jean-Xavier, 2008; Boulenguez et al., 2010). We and others have previously shown that exercise increases KCC2 expression and function after SCI, and that this correlates with improvements in H-reflex RDD (Côté et al., 2014; Chopek et al., 2015; Tashiro et al., 2015). It has also been shown that blockers of KCC2 activity impair the RDD in intact rats (Boulenguez et al., 2010), remove the beneficial effects of exercise in rats with SCI when delivered acutely (Côté et al., 2014) and prevent the beneficial effects of exercise when delivered chronically (Beverungen et al., 2019). In this study, we demonstrate that pharmacologically increasing KCC2 activity with CLP257 in the lumbar spinal cord reduces correlates of spasticity.

### CLP257 increases KCC2 membrane expression on lumbar motoneurons

After chronic SCI, KCC2 expression is decreased on lumbar motoneuronal membranes, while the presence of cytoplasmic clusters increases (Boulenguez et al., 2010; Côté et al., 2014), suggesting changes in post-translational membrane insertion and endocytosis of KCC2 (Lee et al., 2007; Lee et al., 2010). Although the precise mechanism of action remains unknown, CLPs have been shown to increase chloride extrusion by rescuing plasmalemmal KCC2 protein turnover post-translationally (Gagnon et al., 2013; Ferrini et al., 2017). In addition, its effect on locomotion has been replicated by virally overexpressing KCC2 in spinal inhibitory neurons (Chen et al., 2018). While the effectiveness of CLP257 has been challenged (Cardarelli et al., 2017), our results further confirm that CLP257 increases cell surface expression of KCC2 in spinal neurons (Gagnon et al., 2017; Chen et al., 2018).

### Enhancing KCC2 decreases hyperreflexia after chronic SCI

The hyperexcitability of the stretch reflex is a defining hallmark of spasticity (Lance, 1980), and can be assessed either through electrical stimulation of the nerve (H-reflex), or through a mechanical stretch of the muscle. In our experiments, slow stretches of the triceps surae elicited no EMG activity in SCI animals, whereas high velocity stretches evoked both large phasic monosynaptic EMG responses, as well as long lasting tonic activity. While exaggerated monosynaptic reflexes contribute to clonus and hyperreflexia (Bennett et al., 1996; De Serres et al., 2002), the long-lasting responses are thought to contribute to hypertonia and initiation of spasms (Bennett et al., 1999; 2004; Li et al., 2004; Murray et al., 2011b). CLP257 significantly reduced both the phasic and tonic responses, suggesting an effect on multiple spinal pathways that contribute to spasticity (Murray et al., 2010; 2011a; D’Amico et al., 2013a-b; Lucas-Osma et al., 2019). CLP257 did not affect passive force output however, which is not surprising given that passive muscle force is not decreased by SCI in both cats (Frigon et al., 2011) and in humans (Mirbagheri et al., 2001).

CLP257 also improved the H-reflex RDD after chronic SCI, indicating that increasing KCC2 activity restores modulation of the monosynaptic reflex. Yet, while the phasic response to muscle stretch was decreased by CLP257, the basic properties of unconditioned H-reflexes remained unaffected. This is likely because of differences in how responses are generated (Birnbaum and Ashby, 1982; Burke et al., 1984): electrical stimulation directly and synchronously activates group I afferents, while tendon stretch mechanically activates muscle spindles. As a result, mechanical stretch produces a more dispersed afferent volley and longer-lasting activation of the motor pool than electrical stimulation. Thus, responses to a single mechanical stretch are more likely affected by postsynaptic inhibition than H-reflexes, and more likely modulated by KCC2 activity.

Restoring KCC2 membrane expression in lumbar motoneurons following either a complete transection or contusion injury was shown to improve allodynia, neuropathic pain, and locomotion (Boulenguez et al., 2010; Tashiro et al., 2015; Liabeuf et al., 2017; Sanchez-Brualla et al., 2018). Our results show that enhancing KCC2 expression/function in lumbar motoneurons can also ameliorate symptoms of spasticity. While excitability of motoneurons ultimately dictates the final motor output, increasing KCC2 activity in inhibitory relay interneurons has proven to have significant effects in staggered hemi-transected mice. (Chen et al., 2018). While there are significant differences between the mouse and rat model (further discussed in Côté, 2020), this nevertheless suggests that restoring chloride extrusion in spinal interneurons may also contribute to functional recovery. As spinal interneurons play a role in the initiation of spasms (Bellardita et al., 2017; Lin et al., 2019), CLP257 may have affected these neurons in our study as well. Whether this additionally contributed to the observed decrease in the tonic component of the stretch-reflex remains to be determined.

### KCC2 enhancers mimic the effect of exercise on hyperreflexia after chronic SCI

After chronic SCI, rehabilitation programs prevent the downregulation of KCC2, which correlates with improvements in reflex modulation. Similarly, exercised animals in this study displayed increased levels of KCC2 membrane expression in lumbar motoneurons, and had a RDD of the H-reflex that resembled those of intact rats (Thompson et al., 1992; Boulenguez et al., 2010; Côté et al., 2014; Beverungen et al., 2019). In addition, exercised animals displayed no tonic EMG responses to muscle stretches, and smaller phasic responses. In sedentary animals, CLP257 affected both phasic and tonic EMG responses and RDD in a similar manner to exercise. This suggests that KCC2 enhancers mimic the effects of a rehabilitation program on several correlates of spasticity.

Interestingly, CLP257 did not enhance the benefits of exercise on spastic symptoms. However, it did not impede the benefits either. The former provides further evidence that exercise restores reflex modulation via an increase in KCC2 activity; the latter suggests that KCC2 enhancers may be used in tandem with exercise-based therapies without detrimental effects. These results indicate that KCC2 enhancers may be a promising alternative for SCI individuals and could reduce the need for exercise-based rehabilitation and the associated costs.

### KCC2 enhancers increase spontaneous activity in exercised animals without hindering benefits on spasticity

There was a notable presence of spontaneous EMG activity in exercised animals. CLP257 further increased this activity, while blocking KCC2 with VU0240551 returned the activity back to baseline levels. This suggest that while increasing KCC2 activity reduces hyperreflexia, it does not decrease overall spinal excitability and increases activity in certain pathways, perhaps by restoring inhibition on inhibitory interneurons within the spinal cord (Chen et al., 2018). Importantly, the spontaneous activity we observe appears distinct from muscle spasms. It resembles activity often seen in SCI individuals, which involves weak contractions (often unnoticed by subjects) that have no obvious trigger (Zijdewind and Thomas, 2001). Spasms, in contrast, involve strong contractions, vary in intensity and duration, and are easily initiated (Kawamura et al., 1989; Little et al., 1989). In fact, studies have reported spontaneous activity either to have no correlation (Kirshblum et al., 2001) or to be negatively correlated (Campbell et al., 1991) with spasticity in patients. However, the topic is still nebulous. Our results provide an unexpected lead in understanding how this activity relates to spasticity, exercise, and KCC2 function, and merits further investigation in future studies.

### Clinical benefits of increasing KCC2 activity

Current pharmacological treatments for spasticity result in severe side effects, including depression of motor function, seizures, hallucinations, liver toxicity, impaired memory, and possible addiction (Kita and Goodkin, 2000; Burchiel and Hsu, 2001; Elovic, 2001; Adams and Hicks, 2005). All of these severely hinder functional recovery. This work has demonstrated the beneficial effects of pharmacologically increasing KCC2 with CLPs on spastic symptoms after chronic SCI. There are many potential advantages to this strategy. KCC2 is expressed solely in the CNS (Rivera et al., 1999; Medina et al., 2014), so is less likely to produce significant peripheral effects. Additionally, by restoring endogenous inhibition rather than depressing global excitability, restoring chloride homeostasis is likely to avoid detrimental effects on locomotor rehabilitation seen with current antispastic medications. Indeed, when using CLP to treat neuropathic pain in rats, Gagnon et al. (2014) found that CLP had no effect on performance in a rotarod assay, suggesting that CLP has no motor side effects. What’s more, CLPs were recently found to improve spontaneous locomotor recovery in staggered hemi-transected mice (Chen et al., 2018). This, along with our results, implicates pharmacological manipulation of KCC2 as a promising new strategy to alleviate symptoms of spasticity in individuals with SCI.

## Acknowledgements

We thank Drs. Yves de Koninck and Annie Castonguay from Université Laval for the generous gift of CLP257 and technical assistance. This work was supported by grants from the National Institute of Neurological Disorders and Stroke (RO1 NS083666) and the Craig H. Neilsen Foundation (189758).

