## Supplementary material for "Enhancing KCC2 activity decreases hyperreflexia and spasticity after chronic SCI": Table 1

Table 1. Properties of the M-Wave and H-Reflex: Latency, Amplitude, and Threshold

|  | | n= | Motor threshold | M_max_ | H_max_ | M latency | H latency | H-reflex threshold | H_max_ | H_max_/M_max_ |
| --- | --- | --- | --- | --- | --- | --- | --- | --- | --- | --- |
|  | |  | (mA) | (mV) | (mV) | (msec) | (msec) | (x MT) | (xMT) | Ratio |
| SCI | Pre-CLP257 | 10 | 0.031 ± 0.015 | 6.97 ± 1.00 | 2.37 ± 0.40 | 2.63 ± 0.06 | 9.37 ± 0.08 | 1.08 ± 0.07 | 1.46 ± 0.09 | 0.37 ± 0.07 |
|  | Post-CLP257 | 10 | 0.036 ± 0.018 | 6.62 ± 1.22 | 2.18 ± 0.54 | 2.66 ± 0.08 | 9.45 ± 0.12 | 1.09 ± 0.07 | 1.39 ± 0.11 | 0.34 ± 0.05 |
|  | Pre-Saline | 8 | 0.080 ± 0.025 | 5.75 ± 1.27 | 1.20 ± 0.25 | 2.54 ± 0.10 | 8.99 ± 0.22 | 1.09 ± 0.05 | 1.37 ± 0.09 | 0.24 ± 0.04 |
|  | Post-Saline | 8 | 0.083 ± 0.026 | 5.71 ± 1.21 | 1.32 ± 0.27 | 2.47 ± 0.09 | 8.96 ± 0.24 | 1.10 ± 0.05 | 1.38 ± 0.07 | 0.27 ± 0.04 |
| SCI | Pre-CLP257 | 11 110 | 0.032 ± 0.012 | 8.44 ± 1.34 | 3.51 ± 0.87 | 2.48 ± 0.09 | 9.33 ± 0.28 | 0.92 ± 0.09 | 1.20 ± 0.15 | 0.41 ± 0.06 |
| + Ex | Post-CLP257 | 11 | 0.071 ± 0.023 | 7.77 ± 1.03 | 3.96 ± 0.90 | 2.45 ± 0.10 | 9.40 ± 0.25 | 0.99 ± 0.13 | 1.51 ± 0.21 | 0.49 ± 0.08 |
|  | Pre-Saline | 5 | 0.060 ± 0.025 | 7.93 ± 2.58 | 2.50 ± 0.37 | 2.40 ± 0.14 | 9.25 ± 0.25 | 1.28 ± 0.19 | 1.49 ± 0.17 | 0.40 ± 0.09 |
|  | Post-Saline | 5 | 0.067 ± 0.035 | 8.06 ± 1.96 | 3.14 ± 0.39 | 2.44 ± 0.13 | 9.25 ± 0.24 | 1.16 ± 0.12 | 1.44 ± 0.11 | 0.46 ± 0.08 |

Values are mean

±

SEM. MT, motor threshold.
